# Identification of Miro as a mitochondrial receptor for myosin XIX

**DOI:** 10.1101/296376

**Authors:** Stefanie J. Oeding, Katarzyna Majstrowicz, Xiao-Ping Hu, Vera Schwarz, Angelika Freitag, Ulrike Honnert, Petra Nikolaus, Martin Bähler

## Abstract

Mitochondrial distribution in cells is critical for cellular function and proper inheritance during cell division. In mammalian cells, mitochondria are transported predominantly along microtubules by kinesin and dynein and along actin filaments by myosin. Myosin XIX (Myo19) associates with the outer mitochondrial membrane, but no specific receptor has been identified. Using proximity BioID labeling, we identified Miro-1 and Miro-2 as potential binding partners of Myo19. Interaction studies show that Miro-1 binds to a C-terminal fragment of the Myo19 tail region and that Miro recruits the Myo19 tail *in vivo*. This recruitment is regulated by the nucleotide-state of the N-terminal Rho-like GTPase domain of Miro. Notably, Myo19 protein stability in cells depends on its association with Miro. Finally, Myo19 regulates the subcellular distribution of mitochondria. Downregulation, as well as overexpression, of Myo19 induces perinuclear collapse of mitochondria, phenocopying the loss of kinesin KIF5 or its mitochondrial receptor Miro. These results suggest that Miro coordinates microtubule- and actin-based mitochondrial movement.

## INTRODUCTION

Mitochondria are dynamic organelles that can fuse with each other, divide, or move along cytoskeletal tracks, and make functional contacts with other membranous compartments (Friedman and Nunnari, 2014). Interfering with this dynamic behavior impairs mitochondrial function, and may lead to a multitude of mostly degenerative diseases (Nunnari and Suomalainen, 2012; Mishra and Chan, 2014).

In cells, mitochondria are constantly transported to sites where they are needed. This transport occurs either along actin filaments or microtubules depending on the organism. In animal cells, mitochondria are predominately hauled bidirectionally along microtubules. The microtubule-based motor kinesin-1 (KIF5) transports mitochondria towards the plus-end of microtubules that usually points towards the cell periphery (Tanaka et al., 1998). KIF5 is recruited to mitochondria with the help of adaptor proteins (Stowers et al., 2002), which mediate the binding to the outer mitochondrial membrane protein Miro (Miro-1 and Miro-2 in vertebrates). Miro encompasses a C-terminal transmembrane domain and two EF-hand or ELM domains that are in turn flanked by two GTPase domains (Fransson et al., 2003; Klosowiak et al., 2013; see also supplementary figure S1). The N-terminal GTPase domain shares similarity with the Rho-subfamily of monomeric GTPases. An adaptor protein that links KIF5 with Miro was first identified in Drosophila (Glater et al., 2006). Subsequently, two mammalian homologues, TRAK1 (OIP106) and TRAK2 (GRIF-1), were identified as adaptors and shown to mediate bidirectional microtubule-dependent transport of mitochondria (Brickley et al., 2005; Fransson et al., 2006; McAskill et al., 2009; Koutsopoulos et al., 2010; Brickley and Stephenson, 2011; van Spronsen et al., 2013). Transport of mitochondria towards the microtubule minus-ends is mediated by the dynein/dynactin complex. This complex similarly interacts with the adaptors TRAK1 and TRAK2 (van Spronsen et al., 2013; Gama et al., 2017). The microtubule-dependent transport of mitochondria can be arrested by the binding of Ca^2+^ to the EF-hands of Miro (Saotome et al., 2008; MacAskill et al., 2009b; Cai and Sheng, 2009; Wang and Schwarz, 2009). A point mutation in the N-terminal GTPase domain of Miro that is predicted to favor the GDP-bound state, revealed that this domain is essential for mitochondrial transport and morphology (Fransson et al., 2003; Babic et al., 2015). However, it is currently not known whether this GTPase domain cycles between different nucleotide-bound states under physiological conditions. The protein Alex3 of the Armcx genes also regulates mitochondrial movements, in neurons, by an interaction with the Miro/TRAK2 complex and this interaction is abrogated by Ca^2+^-binding to the EF-hands of Miro (Lopéz-Doménech et al., 2012). In addition, Miro can recruit the protein CenpF, and thereby contributes to the mitotic redistribution of mitochondria (Kanfer et al., 2015). The proteins Mitofusin 1/2, DISC1 and APC are yet other proteins that were reported to interact with the Miro/TRAK complex (Misko et al., 2010; Norkett et al., 2015; Mills et al., 2016). The protein hypoxia up-regulated mitochondrial movement regulator (HUMMR) was shown to interact directly with Miro (Li et al., 2009). Importantly, Miro is also a substrate for the E3 ubiquitin ligase Parkin that targets it for proteasomal degradation (Sarraf et al., 2013). Mutations in Parkin have been associated with Parkinson disease (Kitada et al., 1998).

Actin filaments in mammalian cells critically contribute to mitochondrial fission. The ER-associated formin INF2 in combination with a mitochondria-associated splice form of the actin nucleator Spire and the filament forming motor myosin II are involved in constricting mitochondria to initiate their fission (Korobova et al., 2013, 2014; Manor et al., 2015; Ji et al., 2015). Additionally, actin filaments serve as tracks for mitochondrial movement. The actin-based motor myosin XIX (Myo19) is associated with mitochondria and when overexpressed induces the formation of motile tadpole-shaped mitochondria (Quintero et al., 2009). Downregulation of Myo19 interferes with the proper partitioning of mitochondria during mitosis, leading to a stochastic failure in cell division (Rohn et al., 2014). Myo19 consists of a head, light chain binding and tail domain (see supplementary figure S1). *In vitro* studies revealed that the head domain is a plus-end directed motor that is firmly attached to F-actin for most of the time of its chemo-mechanical cycle (Lu et al., 2014). The tail domain, on the other hand, specifies its recruitment to mitochondria (Quintero et al., 2009). A short motif in the tail domain (amino acids 860-890) is able to mediate binding to the lipids of the outer mitochondrial membrane (Shneyer et al., 2016; Hawthorne et al., 2016). However, it is not known how targeting specificity to the outer mitochondrial membrane is achieved and whether there is any coordination between actin- and microtubule-based movements of mitochondria.

In the present study, we attempted to gain further insight into the functions of Myo19 by investigating the molecular mechanism(s) of its recruitment to mitochondria. Interestingly, Myo19 shares its mitochondrial receptor with kinesin (KIF5) and dynein that potentially allows for a coordination of actin- and microtubule-based mitochondrial movements. A preliminary account of this work has been published in the form of an abstract (Mol. Biol. Cell 2016 27:25 3947, *P947*; doi:10.1091/mbc.E16-10-0736).

## RESULTS

### Miro is a mitochondrial receptor of Myo19

To identify potential mitochondrial receptors for Myo19, we used tail proximity-dependent biotinylation (BioID) (Roux et al., 2012), since the tail region contains the mitochondrial targeting information and localizes to mitochondria (Quintero et al., 2009). We generated a stably transfected HeLa cell clone that expressed low levels of the fusion protein BirA*-Myo19 tail when incubated with sodium-butyrate (Fig. 1 A, B). Biotinylated proteins were affinity purified by magnetic streptavidin-beads and analysed by mass spectrometry (Table1). Thereby, we identified peptides from Miro-1 and Miro-2, the mitochondrial receptors for microtubule-based transport. Immunoblotting experiments further demonstrated that Miro, but not the outer mitochondrial membrane protein VDAC-1, was biotinylated specifically by BirA*-Myo19 tail. Miro was eluted after affinity purification from streptavidin exclusively upon expression of the BirA*-Myo19 tail construct (Fig. 1 C, D). In accordance with our results, Miro-2 has recently been identified as a binding partner of Myo19 in two independent large-scale protein-protein interaction screens (Huttlin et al., 2015; Hein et al., 2015). A few additional outer mitochondrial membrane proteins were discovered in the Myo19-tail BioID screen, namely metaxin-3, AKAP1 and MAVS (Table1). Subsequently, we focused on Miro proteins and tested whether Myo19 interacts with Miro1 directly (Fig. 2). We expressed and purified C-terminally His-tagged recombinant human Miro1 lacking its transmembrane domain (Miro1-ABC, aa 1-592; Klosowiak et al., 2016) (Fig. 2A). As a potential binding partner, we expressed and purified a FLAG-tagged C-terminal tail fragment (aa 898-970-FLAG) of human Myo19 that lacks the more N-terminally located lipid-binding motif (Fig. 2A). The purified Myo19 C-tail fragment still contained a major contaminant protein of approx. 70 kD from *E. coli*. Pull-down experiments were performed with Miro1 adsorbed to Talon beads. The Myo19 C-tail fragment was eluted specifically together with Miro, but not from control beads lacking Miro1-ABC (Fig. 2B), indicating that this Myo19 tail fragment interacts directly with Miro1.

**Table 1:**
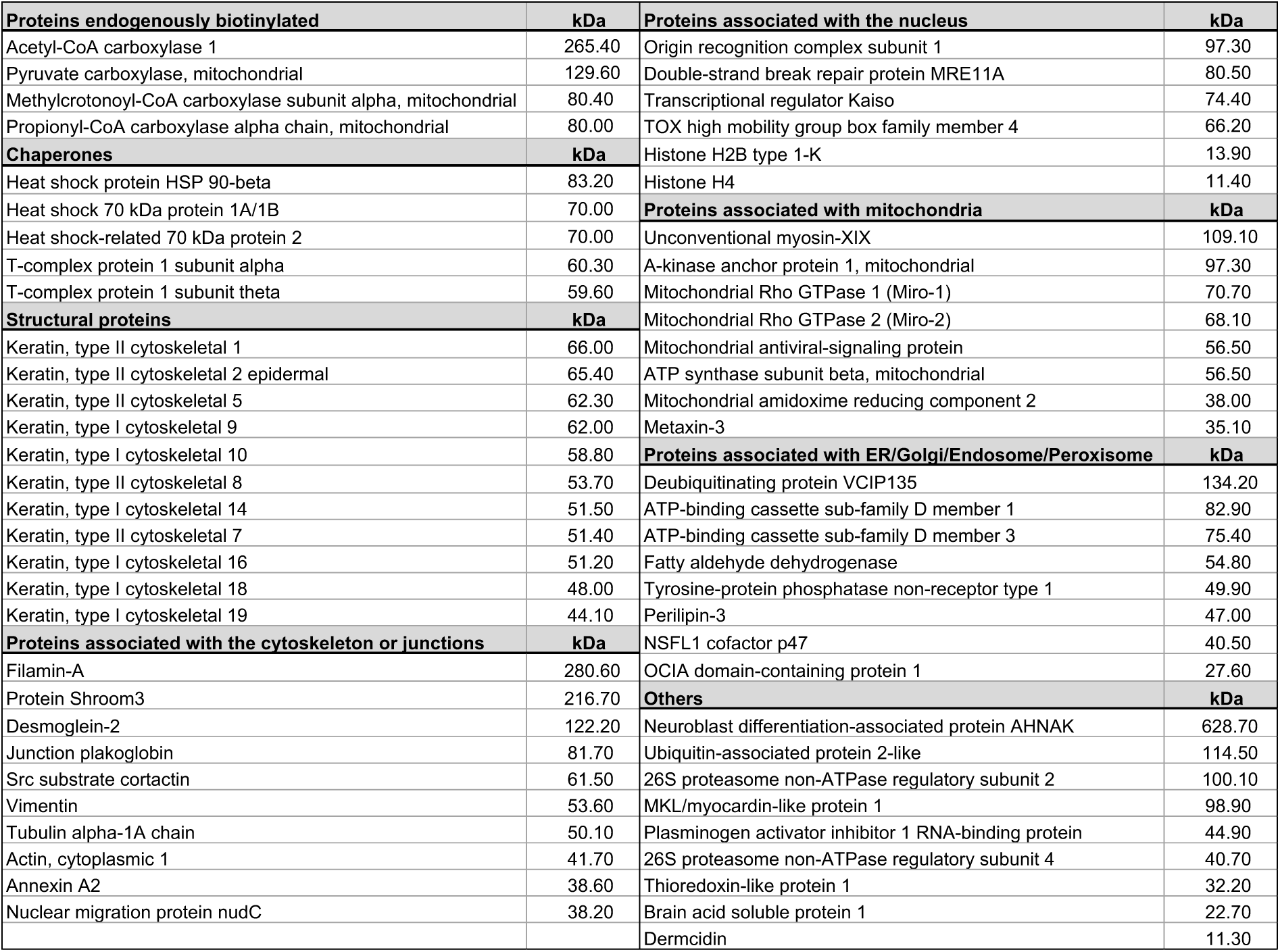
Proteins identified by BioID. Proteins biotinylated by myc-BirA*-Myo19 tail were affinity purified with streptavidin-coupled magnetic beads and their identity was determined by mass spectrometry. Proteins are classified as depicted in the chart and sorted by molecular size. Results are from 3 independent experiments.

**Figure 1:**
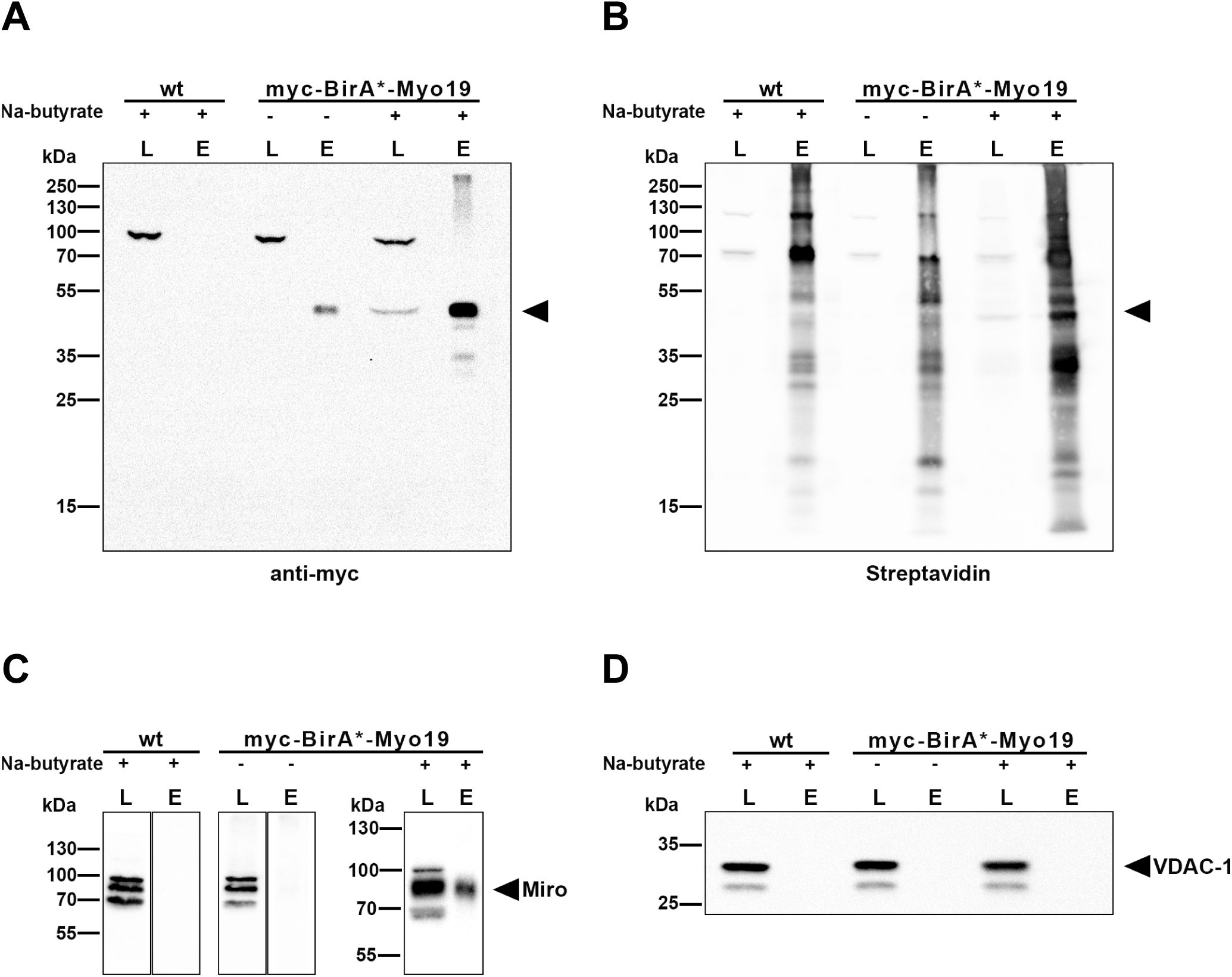
Identification of potential Myo19 tail binding proteins by BioID. Stably transfected myc-BirA*-Myo19 tail HeLa cells and HeLa wild-type (wt) cells were grown in medium supplemented with biotin. Cells were either treated overnight with Na-butyrate (+) or not (-) to enhance the expression of the recombinant construct. Biotinylated proteins were purified by streptavidin affinity-chromatography. Lysates (L) and eluates (E) were separated by SDS-PAGE and transferred to a PVDF membrane for immunoblotting. A) The protein myc-BirA*-Myo19 tail was detected by immunoblotting with anti-myc antibody (arrowhead). Stably transfected cells treated with Na-butyrate showed a much stronger expression of the myc-BirA*-Myo19 tail construct than cells without. B) Immunoblot analysis with peroxidase-labeled streptavidin shows strongly enhanced biotinylation of endogenous proteins after elevated myc-BirA*-Myo19 tail expression in cells treated with Na-butyrate (arrowhead indicates biotinylated construct). C) Biotinylation of Miro, as a function of myc-BirA*-Myo19 tail expression, enhanced by Na-butyrate treatment, is shown by immunoblotting of cell lysates and eluates from streptavidin-beads. The Miro antibody is recognizing Miro-1 and Miro-2 and is cross-reacting nonspecifically in the lysates with proteins running above and below Miro. Cell lysates (L) and eluates (E) from streptavidin-beads arising from wt and stably transfected HeLa cells were loaded on the same gel, but not next to each other. D) The outer mitochondrial membrane porin VDAC-1 did not get biotinylated by myc-BirA*-Myo19 tail.

**Figure 2:**
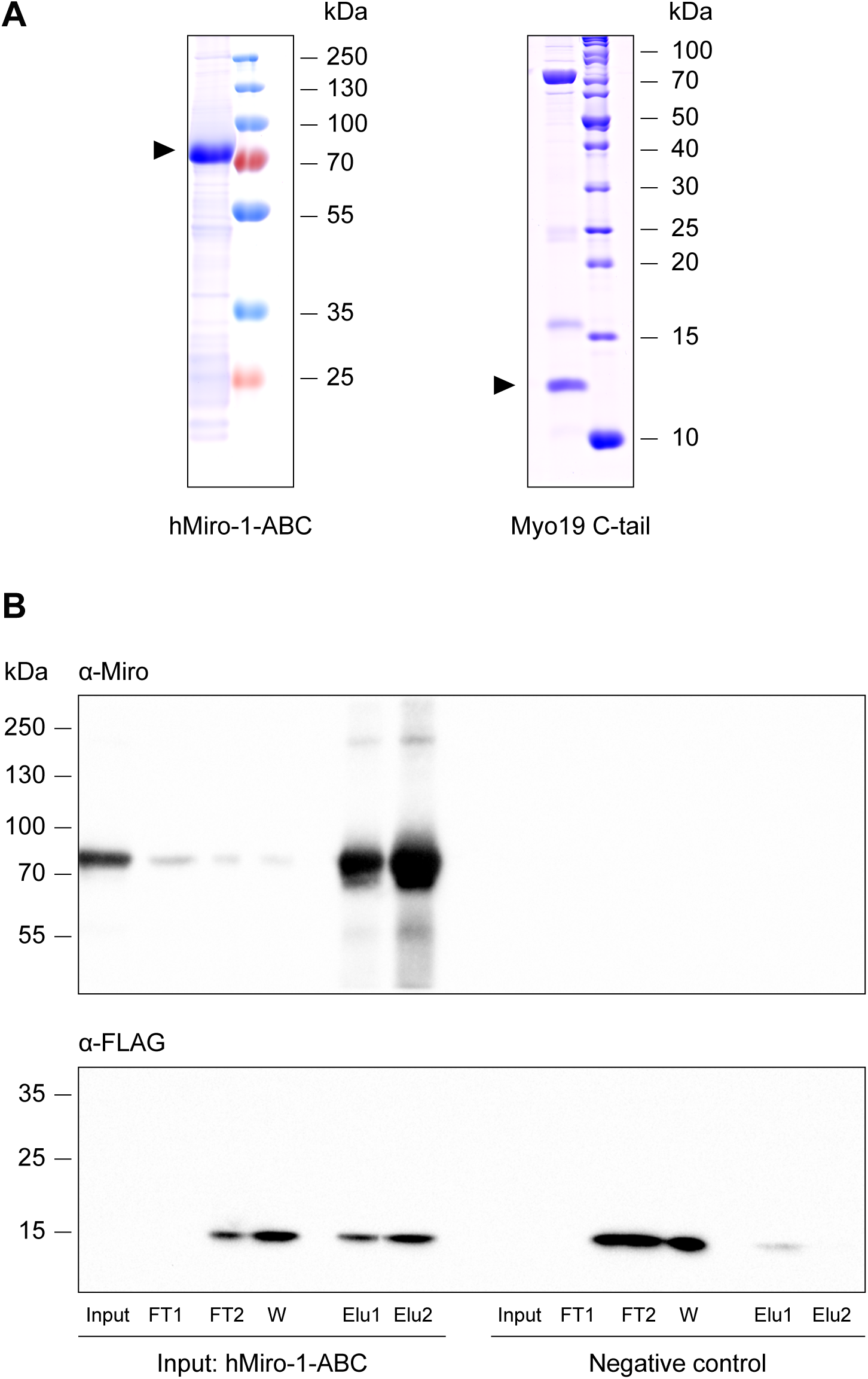
Myo19 C-tail binds directly to Miro-1 in vitro. A) Recombinant Miro1-ABC (aa 1-592) and Myo19 C-tail (aa 898-970) were expressed in *E. coli* and purified as described in Materials and Methods. The purified proteins were separated by SDS-PAGE and stained with Coomassie blue. Miro1-ABC and Myo19 C-tail bands are indicated by arrowheads and molecular mass markers are indicated on the right. B) Pull-down assay showing that Myo19 C-tail binds to His-tagged Miro1-ABC directly. Purified Miro1-ABC (input) was bound to TALON beads and incubated with purified Myo19 C-tail. After washing, Miro1-ABC was eluted with an excess of imidazole. Western blot analysis with antibodies directed against Miro and the FLAG-tag of Myo19 C-tail show coelution of Miro1-ABC and Myo19 C-tail. Input: Purified Miro protein that was bound to TALON beads. FT1: Flow-Through of Miro1-ABC after incubation with TALON beads; FT2: Flow-Through of Myo19 C-tail after incubation with TALON beads. W: Wash. Elu1 + 2: Eluted fractions of Miro1-ABC. As control, beads were incubated with buffer and Myo19 C-tail only. Molecular mass markers are indicated on the left.

To test for potential involvement of Miro in the localization of Myo19 to mitochondria, we downregulated Miro (Miro-1 and Miro-2) by siRNA in HeLa cells. Notably, the downregulation of Miro resulted in a concomitant downregulation of Myo19 as revealed by immunoblotting (Fig. 3A-D). This drop in Myo19 protein was not due to a general reduction in mitochondrial mass, as the signal for the outer mitochondrial membrane protein VDAC-1 was not affected by downregulation of Miro (Fig. 3 A, B). Transfection of cells with either pools or individual siRNAs directed against Miro-1 and Miro-2 led to time-dependent downregulation of Miro, with clear reduction after 48 h and 72 h (approx. 85 % reduction) (Fig. 3 A, B). Notably, the downregulation of Miro was accompanied by comparable time-dependent downregulation of Myo19 (approx. 70 % after 72 h). The concomitant downregulation of Myo19 with Miro could be attenuated by transient overexpression of Miro2 resistant to the siRNA (Fig. Fig. 3 B, D). Conversely, downregulation of Myo19 with siRNA did not alter the expression of Miro (Fig. 3 A, C). These results strongly argue that Miro stabilizes Myo19, but that contrariwise Myo19 does not influence the stability of Miro. Indirect immunofluorescence labeling of Myo19 in cells treated with siRNA against Miro1 and 2 confirmed that the amounts of Myo19 protein associated with mitochondria depended on the expression of Miro (Fig. 3E). The mitochondrial signal of Myo19 was greatly reduced in cells that had Miro downregulated. To verify that Miro indeed regulates Myo19 protein stability, pulse-chase experiments were performed with Halo-tagged Myo19. Stable cell lines expressing Halo-tagged Myo19 were incubated with siRNA directed against Miro and the Halo-tag of Myo19 protein was pulse-labeled with the fluorophore TMR. Labeled Myo19 decayed faster when simultaneously Miro was being downregulated (Fig. 3 F). Interestingly, Myo19 protein turnover was slower when the Halo-tag was fused to the C-terminus (Fig. 3G). This was the case irrespective of whether the cells were treated with non-targeting control siRNA or Miro1 and 2 siRNA. These data reveal that the positioning of the tag influences the life-time of Myo19 protein and that Miro increases the life-time of Myo19 protein.

**Figure 3:**
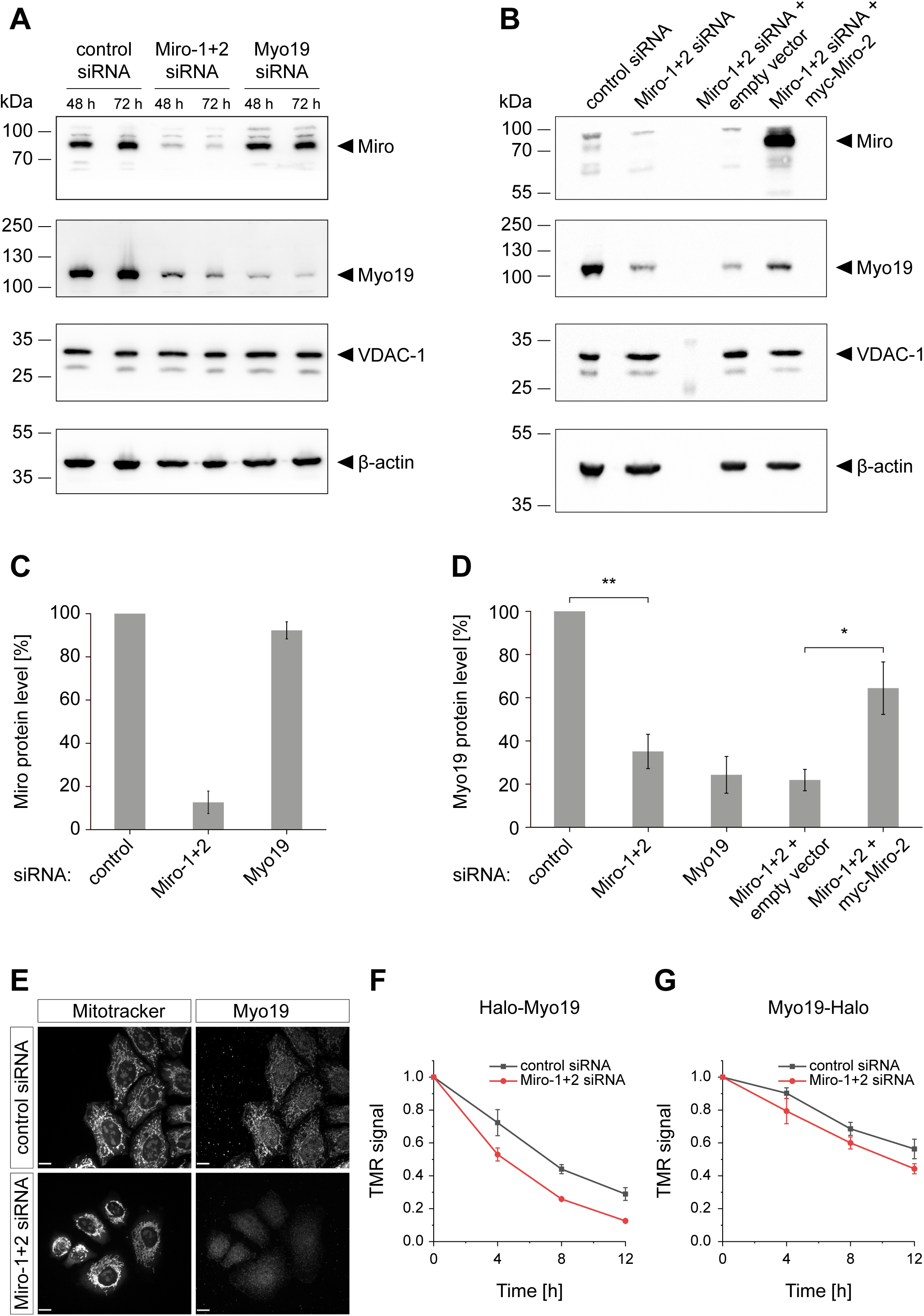
The downregulation of Miro protein concomitantly reduces the amount of Myo19 protein. HeLa cells were transfected with an siRNA pool against Myo19 or cotransfected with a pool of siRNA against Miro-1 and either a pool of siRNA against Miro-2 (A, E) or a specific siRNA against Miro2 (B-D) and the downregulation of Miro and Myo19 was monitored 48 h or 72 h later by immunoblotting with anti-Miro and anti-Myo19 antibodies. Notably, Myo19 protein was downregulated concomitantly with Miro, but not vice versa. Miro knock-down cells were transiently transfected with either empty or myc-Miro2 vector. VDAC-1 and β-actin served as loading controls for equal amounts of mitochondrial mass and total protein, respectively. The experiment was replicated several times with comparable results. C, D) Quantification of the results shown in A) and B). Expression levels of Miro (C) and Myo19 (D) were quantified after the indicated treatments of cells as described in B) from 3 independent experiments. Re-expression of Miro2 in Miro knock-down cells restores Myo19 protein levels. E) Immunofluorescence staining of cells treated with non-targeting or Miro1 and 2 siRNA’s for Myo19. Mitochondria were labeled with Mitotracker. F, G) Pulse-chase labeling of Halo-Myo19 (F) or Myo19-Halo protein (G), respectively, in HeLa cells that were treated with either control siRNA or Miro1 and 2 siRNA. HeLa cells stably expressing Halo-tagged Myo19 were incubated with fluorescent HaloTag-TMR (tetramethylrhodamine) for 30 min. After different times of chase the amount of fluorescent Halo-tagged Myo19 was quantified and plotted.

Further evidence that release of Myo19 from mitochondria promotes its degradation was provided by the observation that stable expression of BirA*-Myo19 tail displaced endogenous Myo19 from mitochondria without corresponding increase of Myo19 protein in the cytosol fraction (Fig. 4). Analysis of cell homogenates confirmed that Myo19 gets degraded upon overexpression of Myo19 tail (Suppl. Fig. S2). Additionally, the overexpression of BirA*-Myo19 tail led to a slight reduction of the amount of TRAK1 that was associated with purified mitochondria. However, the reduction of this adaptor molecule for KIF5B and dynein was not significant (Fig. 4).

**Figure 4:**
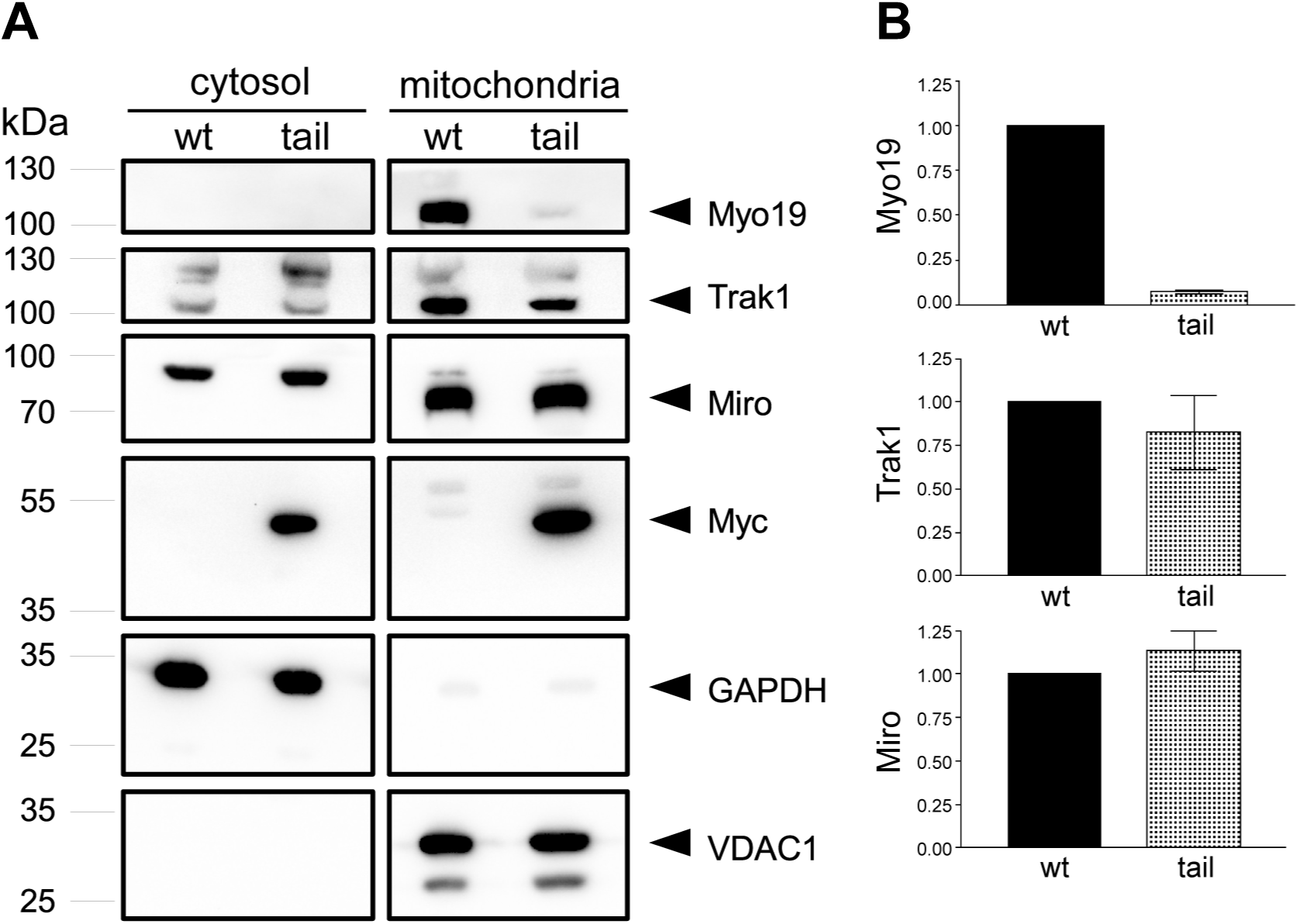
Overexpression of Myo19 tail induces the loss of endogenous Myo19. HeLa wild-type (wt) and stably transfected myc-BirA*-Myo19 tail (tail) cells were incubated overnight with Na-butyrate to enhance the expression of the recombinant construct. Cells were lysed and separated into cytosolic and mitochondrial fractions. A) Immunoblots from a representative experiment are shown. Myo19 is strongly reduced in the mitochondrial fraction of myc-BirA*-Myo19 tail expressing cells and it does not become detectable in the cytosol. GAPDH served as a cytosolic and VDAC1 as a mitochondrial marker, respectively. B) Quantification of band intensities for Myo19, TRAK1 and Miro in the mitochondrial fraction. Band intensities in the mitochondrial fractions were normalized to wt (black bars) and the fold changes are shown for the tail expressing cells. Data are represented as mean ±SEM, n=3.

The simultaneous downregulation of Myo19 with Miro precluded a meaningful analysis of potential redistribution of Myo19 in the absence of Miro. The proportion of background signal significantly increased in relation to the specific signal, so that no meaningful quantification of the remaining mitochondrial localization of Myo19 was possible (see Fig. 3E). Therefore, we devised an alternative strategy to test for *in vivo* recruitment of Myo19 by Miro. We retargeted Miro 1 to early endosomes by replacing its C-terminal transmembrane domain with 2xFYVE domains that mediate binding to phosphatidylinositol-3-phosphate (see schematic representation suppl. Fig. S1). This Miro construct was indeed targeted to early endosomes as it colocalized with fluorescently labeled transferrin that was taken up by receptor-mediated endocytosis (Suppl. Fig. S3A). To monitor retargeting of the Myo19 tail that directs Myo19 to mitochondria (Quintero et al., 2009; see also Fig. 4) and to delineate the mitochondrial and Miro binding site(s) in the Myo19 tail region more precisely, we analysed various fragments of the Myo19 tail region for Miro-dependent retargeting to early endosomes. We found that the tail region of Myo19 localized both to mitochondria with endogenous Miro and to early endosomes with retargeted Miro-1 (Fig. 5 A-C), indicating that Miro is sufficient for the specific recruitment of Myo19 tail *in vivo*. An N-terminal tail fragment (aa 824-897) encompassing a reported lipid-binding domain (Shneyer et al., 2016; Hawthorne et al., 2016) localized exclusively to mitochondria and was not redirected to early endosomes (Fig. 5 A-C). In contrast, the C-terminal tail fragment (aa 898-970) that bound to Miro-1 *in vitro* colocalized with Miro that was retargeted to early endosomes (Fig. 5 A-C). Part of this C-terminal tail fragment was also found in the cytosol, but it was not obviously colocalized with mitochondria (Fig. 5 A-C). These results suggest that the tail region of Myo19 can be targeted to mitochondria by an N-terminal lipid-binding motif and a C-terminal Miro-binding domain. For recruitment of the Myo19 tail region, a fragment of Miro containing the N-terminal GTPase domain and the two ELM domains (aa 1-408) was sufficient (Fig. 5 D-F). The C-terminal GTPase domain was dispensable. Next, we examined potential regulation of Myo19 recruitment by the nucleotide state of the N-terminal GTPase domain of Miro. The introduction of a point mutation (N18) in the N-terminal GTPase domain, which in analogy to other GTPases is predicted to induce a constitutively GDP-bound or nucleotide-free state, abrogated the recruitment of the Myo19 tail and C-terminal tail fragment to mitochondria (Fig. 5 G-I). The correlation between the localization of retargeted Miro and TOMM20 was hardly altered by the different Myo19 tail constructs (Suppl. Fig. S2B).

**Figure 5:**
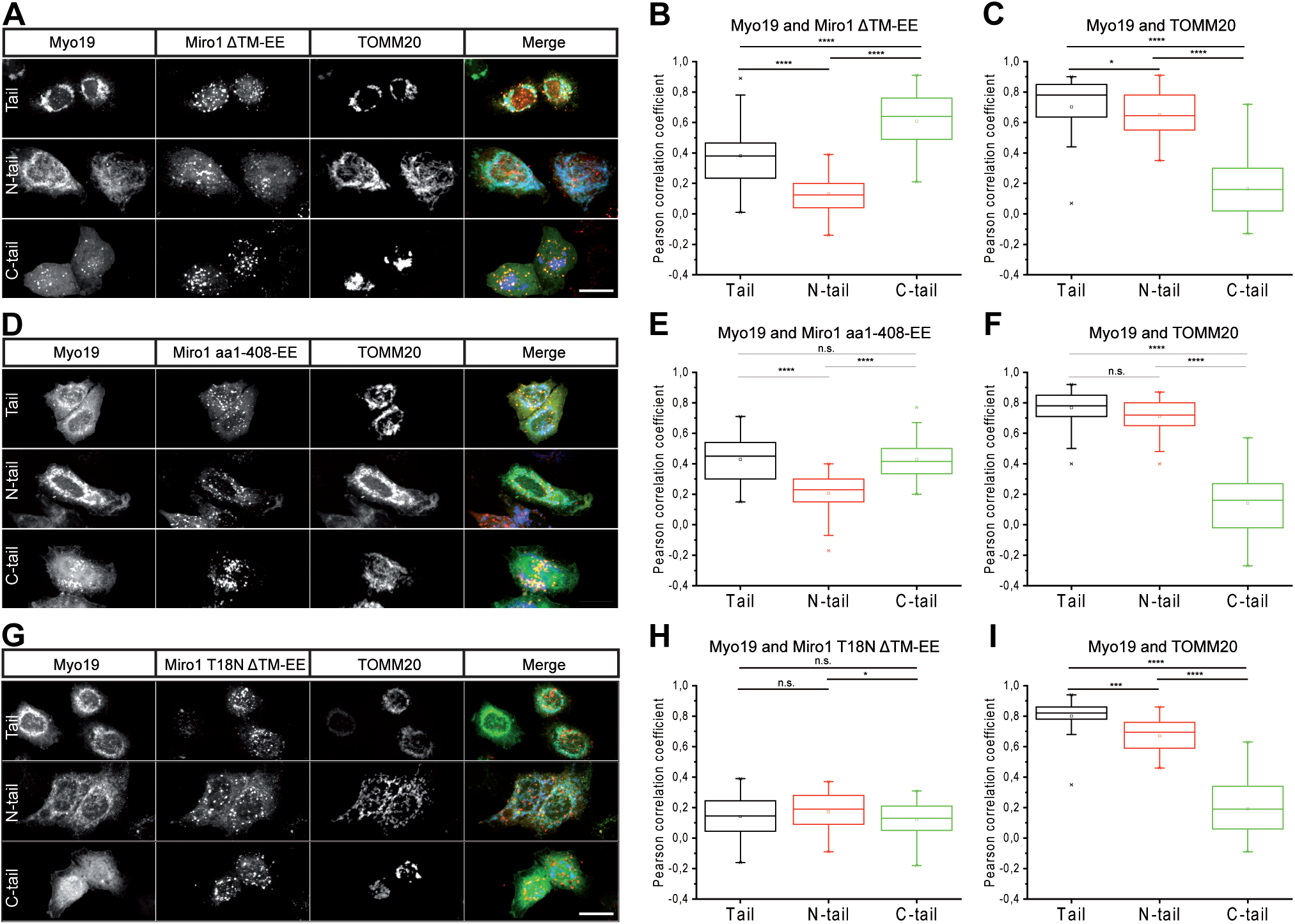
The Myo19 tail contains a Miro-independent and a Miro-dependent mitochondrial targeting site. A) HeLa cells were transfected with EGFP-Myo19-tail (aa 824-970), EGFP-Myo19 N-tail (aa 824-897) or EGFP-Myo19 Myo19 C-tail (aa 898-970) constructs together with mTagBFP2-TOMM20-N-10 and mCherry-MiroΔTM-EE (2xFYVE early endosome targeting sequence (EE)) as indicated. Merged images are shown in color on the right. Scale bar, 20 µm. B,C) Statistical analysis of the correlation between the localization of Myo19 tail constructs (Tail, N-tail, C-tail) with Miro1ΔTM-EE (B) and TOMM20 (C), respectively. Boxplots are shown. The square indicates the sample mean, the crosses the 1^st^ and 99^th^ maximum values, the dashes the minimum and maximum values, respectively. (n≥30 from three independent experiments; ***p ≤ 0.001. n.s.= not significant). D-F) Amino acid residues 1-408 of Miro-1 are sufficient to recruit the Myo19 tail and C-tail constructs. D) HeLa cells were transfected with EGFP-Myo19-tail (aa 824-970), EGFP-Myo19 N-tail (aa 824-897) or EGFP-Myo19 C-tail (aa 898-970) constructs together with mTagBFP2-TOMM20-N-10 and mCherry-Miro1-aa1-408-EE as indicated. Merged images are shown in color on the right. Scale bar, 20 µm. E, F) Statistical analysis of the correlation between the localizations of Myo19 tail constructs (Tail, N-tail, C-tail) with Miro1-aa1-408-EE (E) and TOMM20 (F), respectively. Boxplots are shown. The square indicates the sample mean, the crosses the 1^st^ and 99^th^ maximum values, the dashes the minimum and maximum values, respectively. (n≥30 from three independent experiments; ***p ≤ 0.001. n.s.= not significant). G-I) The nucleotide state of the N-terminal GTPase domain of Miro regulates the mitochondrial recruitment of the Myo19 tail and C-tail constructs. G) HeLa cells were transfected with EGFP-Myo19-tail (aa 824-970), EGFP-Myo19 N-tail (aa 824-897) or EGFP-Myo19 C-tail (aa 898-970) constructs together with mTagBFP2-TOMM20-N-10 and the dominant negative mCherry-Miro1(T18N)ΔTM-EE as indicated. Merged images are shown on the right. Scale bar, 20 µm. H, I) Statistical analysis of the correlation between the localizations of Myo19 tail constructs with Miro1(T18N)ΔTM-EE (H) and TOMM20 (I), respectively. Boxplots are shown. The square indicates the sample mean, the crosses the 1^st^ and 99^th^ maximum values, the dashes the minimum and maximum values, respectively. (n≥30; ***p ≤ 0.001. n.s.= not significant).

### Myo19 regulates the subcellular distribution of mitochondria

In mammalian cells, the subcellular distribution of mitochondria is regulated by microtubule-based motor proteins and their mitochondrial receptor Miro (Maeder et al., 2014; Mishra et al., 2016). The silencing of either Miro or Kif5 protein expression has been shown to induce perinuclear accumulation of mitochondria (Tanaka et al., 1998; Liu et al., 2012). To analyze whether Myo19 contributes to mitochondrial distribution in cells, we either silenced or overexpressed Myo19 in HeLa cells. We specifically measured the perinuclear accumulation factor of mitochondria in cells, which we defined as the percentage of mitochondria within the perinuclear region, divided by the fraction of the cell area occupied by this region. This quantification confirmed that depletion of Miro1+2 or Kif5B with siRNAs causes significant perinuclear accumulation of mitochondria (Fig. 6). Interestingly, siRNA-mediated depletion of Myo19 expression also significantly shifted the localization of mitochondria to the perinuclear region (Fig. 6). Next, we addressed if overexpression of Myo19 or different tail fragments would affect mitochondrial distribution. We found that overexpression of GFP-Myo19 phenocopied the effect of Myo19 downregulation (Fig. 7). This effect was independent of the actin-based motor capacity of Myo19, since the GFP-labeled tail region of Myo19, when overexpressed, induced an even more pronounced perinuclear accumulation of mitochondria (Fig. 7). Overexpression of the Myo19 C-tail fragment that has been shown to interact with Miro also induced perinuclear accumulation of mitochondria (Fig. 7). The N-tail lipid-binding fragment might still compete to some extent for binding of endogenous Myo19 to mitochondria, as its overexpression caused a less pronounced, but still significant, perinuclear accumulation of mitochondria (Fig. 7). To exclude that the perinuclear accumulation of mitochondria was due to oligomerization of the fused EGFP label, the experiments were repeated with full-length Myo19 and Myo19 tail, respectively, that were fused with a Halo tag and labeled with TMR. These experiments confirmed that the perinuclear accumulation of mitochondria occurred independently of EGFP, and again showed that the tail region of Myo19 has a stronger effect on mitochondrial distribution than full-length Myo19 (Fig. 8). We also assessed whether overexpression of Myo19 or tail fragments thereof led to a change in cell size or shape. Such changes might impact the quantification of perinuclear clustering. We found that cells overexpressing full-length Myo19 covered a smaller area, had a shorter perimeter and increased circularity (Suppl. Fig. S4). However, this effect seemed to be dependent on the motor domain of Myo19, since none of the Myo19 tail fragments caused similar changes when overexpressed (Suppl. Fig. S4). In cells overexpressing EGFP-tail, for which the most pronounced perinuclear accumulation of mitochondria had been observed, the perinuclear accumulation factors were only weakly correlated to the cell size and shape as assessed by Pearson’s correlation coefficient. The strongest correlation was found between a cell’s perinuclear accumulation factor and its covered area or perimeter (R = 0.46 and 0.43, respectively). This positive correlation rather confirmed than invalidated the observed increased perinuclear accumulation of mitochondria in the smaller Myo19-overexpressing cells, since increased perinuclear clustering in smaller cells would actually be detected less readily. In summary, both the downregulation of Myo19 and the overexpression of full-length Myo19 or Myo19 tail fragments that contain the Miro-binding site induced perinuclear clustering of mitochondria.

**Fig. 6:**
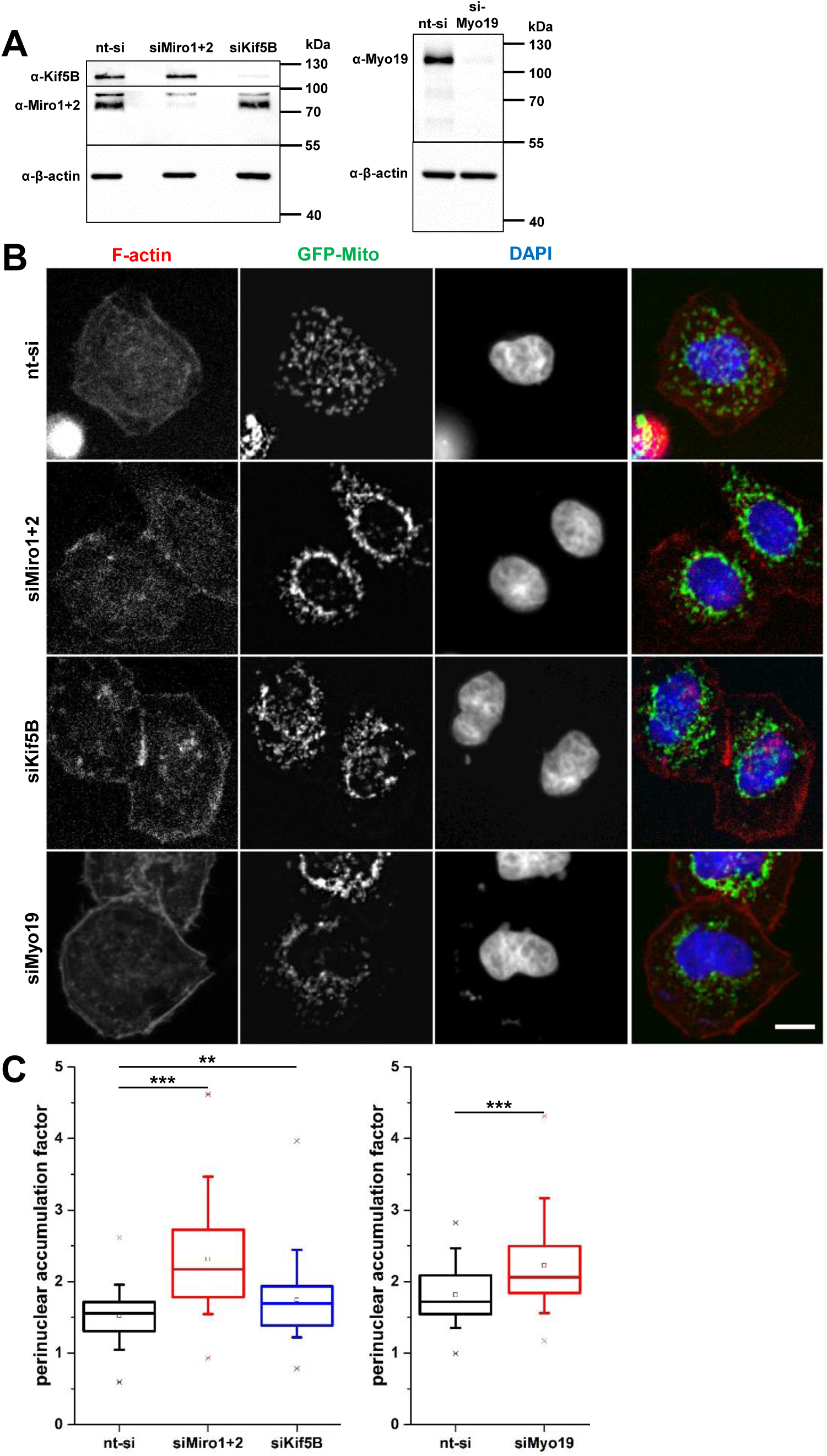
Myo19, Miro1+2 and Kif5B regulate the subcellular distribution of mitochondria in HeLa cells. A) Western blots showing the depletion of Miro1+2 and Kif5B (left panel) and Myo19 (right panel), respectively, 72 h after siRNA treatment as compared to cells treated with nt-siRNA. In these experiments, the reduction of Miro1+2 was 95%, the reduction of Kif5B was 96%, and the reduction of Myo19 was 97%. β-actin served as loading control for equal amounts of total protein. Comparable results were obtained in all mitochondria distribution experiments involving the depletion of Miro1+2, Kif5B or Myo19. B) HeLa cells were transfected with (from top to bottom) nt-siRNA as controls or with siRNAs against Miro1+2, Kif5B or Myo19. Overexpressed GFP-Mito labels mitochondria, F-actin was stained with TexasRed-phalloidin, and nuclei were stained with DAPI. nt-siRNA transfected control cells exhibited normal distribution of mitochondria, whereas depletion of Miro1+2, Kif5B or Myo19 led to clustering of the mitochondria around the nucleus. Scale bar: 10 µm. C) The observed perinuclear accumulation of mitochondria due to depletion of Miro1+2, Kif5B or Myo19 was significant, as measured by the perinuclear accumulation factor. n = 94-97 cells per construct from 3 independent experiments.

**Fig. 7:**
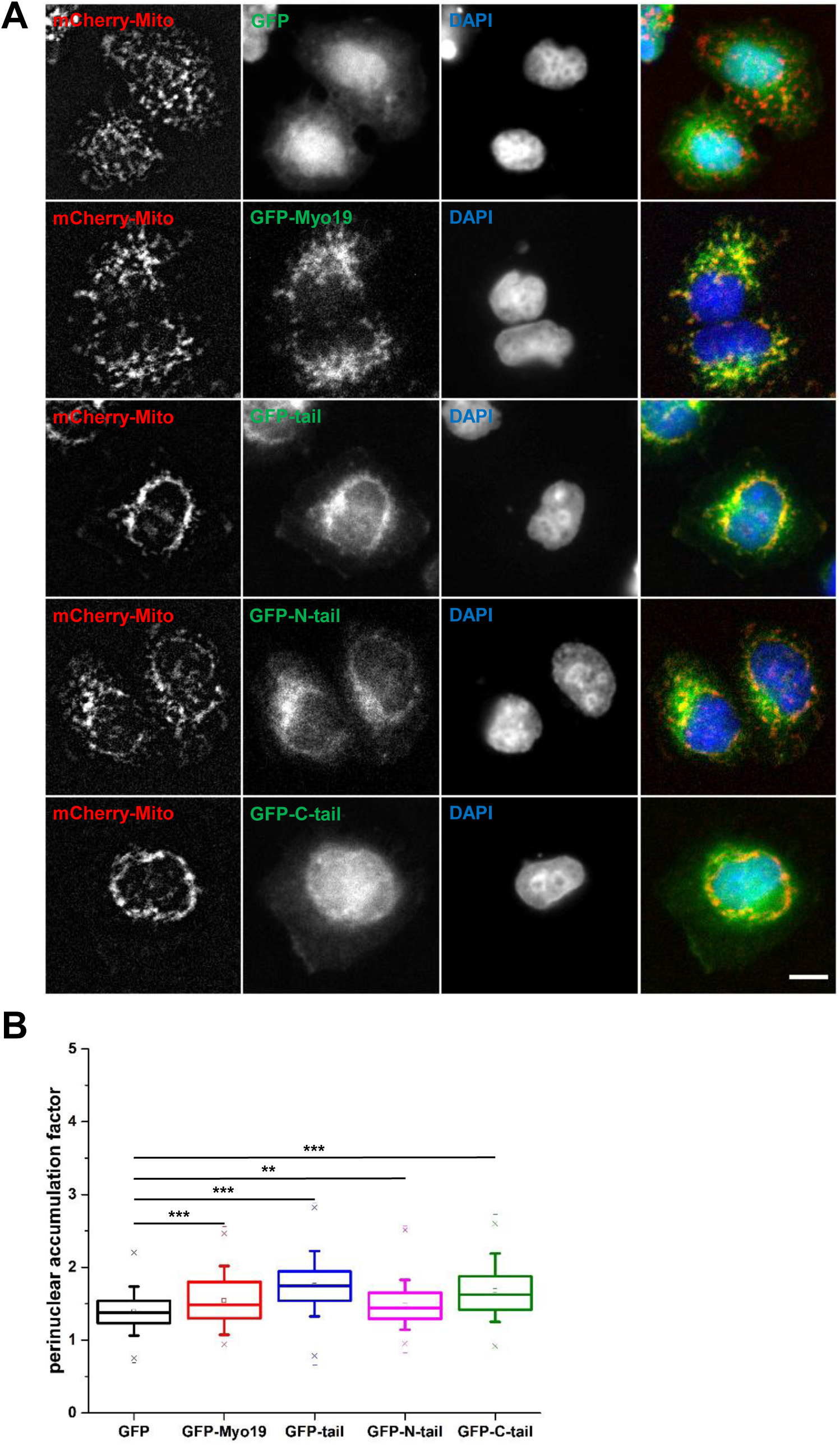
Overexpression of GFP-labelled Myo19 or tail fragments thereof causes perinuclear accumulation of mitochondria in HeLa cells. A) HeLa cells were cotransfected with either (from top to bottom) GFP, GFP-Myo19, GFP-Myo19-tail, GFP-Myo19-C-tail, or GFP-Myo19-N-tail and mCherry-Mito to label mitochondria. Nuclei were stained with DAPI. GFP-overexpressing control cells exhibited normal distribution of mitochondria, whereas the mitochondria were clustered around the nucleus in cells overexpressing Myo19 or tail fragments thereof. The effect is most pronounced for cells overexpressing GFP-My19-tail or GFP-Myo19-C-tail. Scale bar: 10 µm. B) The perinuclear accumulation factor was defined as the percentage of mitochondria within a 4 µm wide region around the nucleus, divided by the percentage of cell area occupied by this region. The overexpression of any of the GFP-Myo19 constructs increased the perinuclear accumulation of mitochondria significantly. n = 153-159 cells per construct from 3 independent experiments.

**Fig. 8:**
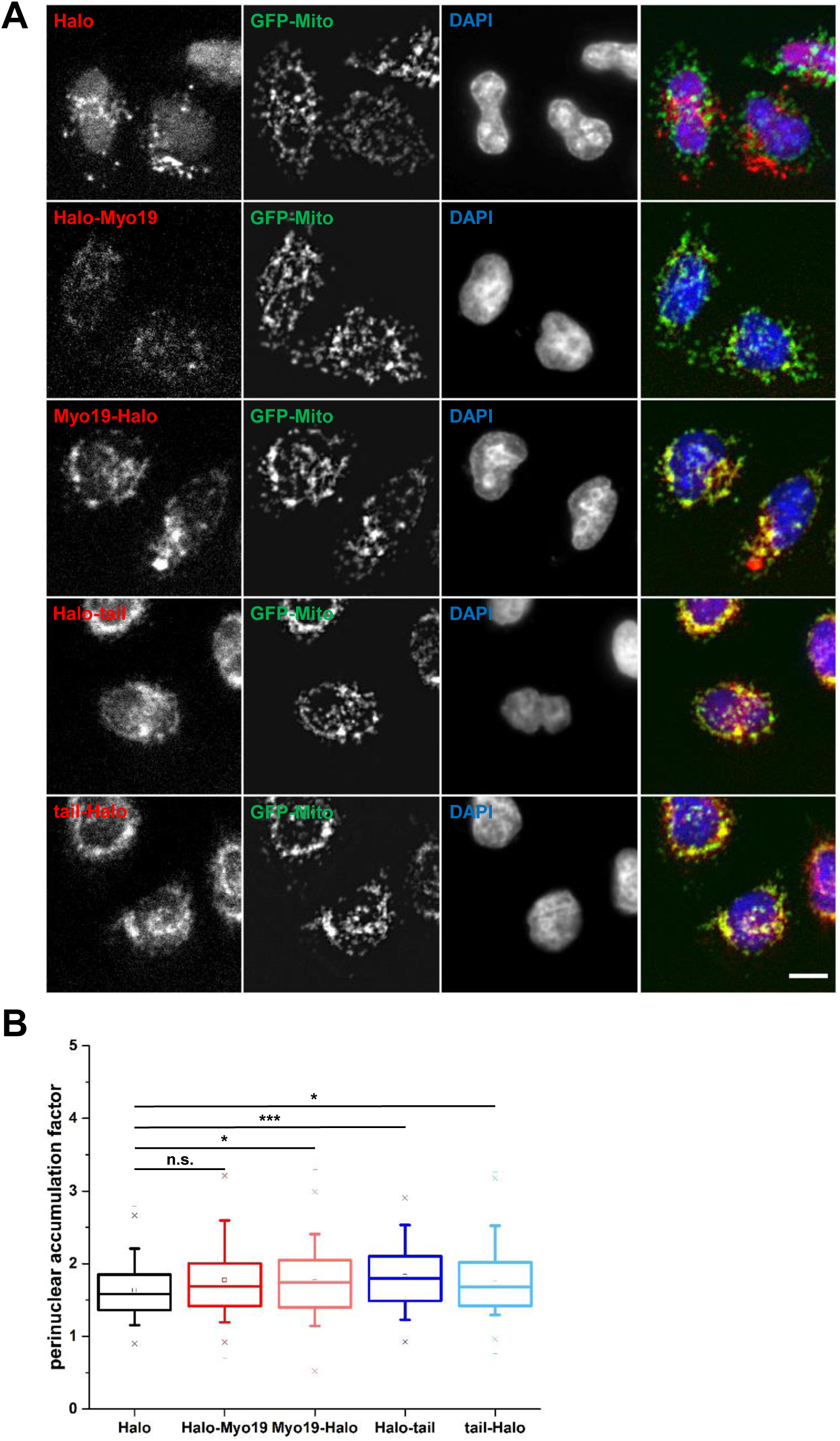
Perinuclear accumulation of mitochondria following overexpression of Myo19 or Myo19 tail fragments is independent of EGFP. A) HeLa cells overexpressing (from top to bottom) Halo, Halo-Myo19, Myo19-Halo, Halo-Myo19-tail, or Myo19-tail-Halo are shown. Mitochondria were labeled by overexpressed GFP-Mito. Halo was labeled with a TMR-Halo ligand and nuclei were stained with DAPI. Halo-overexpressing control cells exhibited a normal distribution of mitochondria, whereas the mitochondria were clustered around the nucleus in cells overexpressing Halo-Myo19-tail or Myo19-tail-Halo. Cells overexpressing full length Halo-Myo19 or Myo19-Halo showed only weak clustering of the mitochondria around the nucleus. Scale bar: 10 µm. B) Overexpression of Halo-Myo19-tail or Myo19-tail-Halo increased the perinuclear accumulation of mitochondria significantly, although the effect was more pronounced for Halo-Myo19-tail. A similar significance was observed for full length Myo19-Halo, but not for Halo-Myo19. n = 114-120 cells per construct from 3 independent experiments.

## DISCUSSION

In this study we identified Miro (comprising Miro-1 and Miro-2) as a mitochondrial receptor for Myo19 and showed that this interaction stabilizes Myo19, but not Miro. We further found that both downregulation of Myo19 and overexpression of Myo19 or Miro-binding Myo19 fragments phenocopy the loss of both Miro and the microtubule-based motor KIF5. KIF5-dependent transport of mitochondria towards the plus-end of microtubules and dynein-dependent transport to the opposite end are mediated by the protein Milton/TRAK that serves as an adaptor for Miro and both KIF5 and dynactin/dynein (Glater et al., 2006; van Spronsen et al., 2013). Although we could demonstrate a direct interaction between Myo19 and Miro, it remains to be determined whether TRAK or another adaptor protein modulates this interaction. This will be an important issue for the future to unravel the coordination between microtubule- and actin-based mitochondrial movements. The perinuclear collapse of mitochondria observed upon both deletion and overexpression of Myo19 could be induced by an imbalance of Miro-mediated microtubule-dependent transport. Deletion of Myo19 might affect the equilibrium of kinesin- and dynein-dependent mitochondrial movements, whereas overexpression might compete differentially with these two motors. Similar to the deletion of Myo19, deletion of Miro caused perinuclear accumulation of mitochondria. The search for nearest neighbors of the Myo19 tail region using BioID did not reveal any known binding partners of Miro. However, additional proteins were identified that are associated with the mitochondrial outer membrane, such as AKAP1 (A-kinase anchor protein 1), MAVS (mitochondrial anti-viral signaling protein) and metaxin-3. The connection of these proteins to Myo19 should be investigated further.

Intriguingly, downregulation of the Myo19 receptor Miro caused concomitant reduction of Myo19 protein. The binding of Myo19 to Miro on mitochondria stabilizes the protein and the loss of this interaction leads to Myo19 destabilization and degradation. Miro can be downregulated by the two Parkinson’s disease genes PINK1 and Parkin that participate in mitochondrial quality control (Chan et al., 2011; Wang et al., 2011). Additionally, the degradation of Miro-2 is regulated by the PGAM5-KEAP1-Nrf2 complex. Miro-2, but not Miro-1, is a substrate for KEAP1-cullin-3 E3 ubiquitin ligase and the proteasome (O’Mealey et al., 2017). Following Miro degradation by either pathway, microtubule-based mitochondrial movement is halted (Wang et al., 2011; O’Mealey et al., 2017). Similarly, actin-based motility by Myo19 will be abrogated by loss of Miro. In agreement with our work, Lopez-Domenech et al. (2018) reported, while this work was in revision that Miro stabilizes Myo19 on mitochondria. A remarkable analogy represents vacuole inheritance in *S. cerevisiae* that relies on actin-based transport by the class V myosin Myo2. Unloading of the vacuole at its proper destination from Myo2 is controlled by the degradation of its vacuole adapter Vac17 (Yau et al., 2017).

Recently, it has been reported that a peptide encompassing residues 860-890 in the tail region of Myo19 serves as an outer mitochondrial membrane-binding motif (Shneyer et al., 2016; Hawthorne et al., 2016). This finding agrees well with our observation that a N-terminal tail fragment (aa 824-897) of Myo19 is able to localize to mitochondria. However, this lipid-binding motif alone appears not to provide sufficient specificity and functionality under physiological conditions. To establish specificity and fully functional association with mitochondria, Myo19 has to interact additionally with Miro. Although the overexpressed Myo19 Miro-binding motif appeared to be localized mainly in the cytosol, it induced perinuclear collapse of the mitochondria, indicating that it successfully competed with other proteins for binding to Miro. An increase of the Miro concentration resulted in the distinct recruitment of the Miro binding C-terminal tail fragment (aa 898-970) of Myo19 to mitochondria (data not shown). Miro was not only necessary, but also sufficient to recruit the Myo19 tail. When Miro was redirected to early endosomes, the Myo19 tail was also localized on early endosomes. Furthermore, this result demonstrates the predominance of the interaction with Miro over the lipid binding by Myo19.

We showed here that the N-terminal Rho-like GTPase domain of Miro is critically involved in the recruitment of Myo19. Currently, there is no conclusive evidence available as to whether this domain cycles between a GDP- and a GTP-bound state *in vivo*. Based on sequence comparisons, it appears likely that this domain in Miro has GTP bound constitutively, and is not cycling, since some residues important for hydrolysis are not conserved (Longenecker et al., 2003; Fransson et al., 2003; Li et al., 2002; Foster et al., 1996; Nobes et al., 1998). Marginal GTPase activity has been reported for this domain in Gem1p, the yeast analog, with one GTP hydrolysed in 5-10 min (Koshiba et al., 2011). Phosphorylation of a serine residue was recently suggested to increase GTPase activity (Lee et al., 2016). Mutation in Miro of a threonine residue at the end of the P-loop that is conserved in nucleotide binding proteins to asparagine (T18N) abrogated the recruitment of Myo19. Introduction of the analogous mutation (T19N) in RhoA produces a dominant negative form. This RhoA mutant has an accelerated rate of GTP hydrolysis, but is mostly in a nucleotide free state exhibiting a high affinity for guanine nucleotide exchange factors (Strassheim et al., 2000; Miyazaki et al., 2002). Genetic experiments in Drosophila with this Miro mutant suggested that it is essentially a recessive loss-of-function mutation (Babic et al., 2015). Residues mutated in dominant active constructs of RhoA are not conserved in Miro-1. Therefore, it is not certain whether the corresponding mutations in Miro-1 will induce similar alterations. Neuronal phenotypes observed for one such point mutation in *Drosophila* (dMiroA20V) were not compatible with a constitutively active mutation (Babic et al., 2015). It remains to be determined whether cycling between different nucleotide states of the N-terminal GTPase domain of Miro represents a physiological mechanism to regulate Myo19 recruitment. The protein VopE from the bacterial pathogen *Vibrio cholerae* has been proposed to act as a GTPase-activating protein (GAP) for the N-terminal GTPase domain of Miro and the mammalian protein RAP1GDS1 as a guanine-nucleotide exchange factor (GEF) (Ding et al., 2016).

Myo19 was shown to regulate the subcellular distribution of mitochondria. Both, downregulation and overexpression of Myo19 induced a perinuclear accumulation of mitochondria. A similar perinuclear accumulation of mitochondria was observed in the absence of Miro which negatively affected cytoplasmic energy distribution, cell migration and cell adhesion (Schuler et al., 2017). During cell division, Myo19 regulates the allocation of mitochondria to daughter cells. Loss of Myo19 causes asymmetric partitioning of the mitochondria to one or both spindle poles in mitosis and stochastic failure of cytokinesis (Rohn et al., 2014). In contrast, in the presence of Myo19, mitochondria do not localize to spindle microtubules, but instead are localized in the periphery (Lee et al., 2007; Chung et al., 2016). Forced attachment of dynein or kinesin motors to the mitochondrial surface during mitosis generates a phenotype comparable to that observed after downregulation of Myo19 (Chung et al., 2016; Rohn et al., 2014). These findings hint at a role for Myo19 in the regulation of microtubule-based transport of mitochondria. Therefore, it will be interesting to elucidate further the interplay between Miro and Myo19 in the control of microtubule- and actin-based transport of mitochondria.

## MATERIALS AND METHODS

### Reagents and antibodies

The antibodies used were as follows: Myo19 (EPR12551-13/ab174286, Abcam, WB: 1:1000; HPA021415, Sigma-Aldrich); Miro-1/2 (NBP1-59021, Novus Biologicals, WB: 1:4000); KIF5B (ab167429; Abcam, WB: 1:1000); TRAK1 (PA5-44180, Thermo Fisher, WB: 1:1000); VDAC1 (20B12AF2/ab14734, Abcam, WB: 1:1000); myc (ab9106, Abcam, WB: 1: 25,000); GAPDH (TA802519; OriGene); β-actin (AC-15/A1978, Sigma-Aldrich, WB: 1:2000); Goat Anti-Rabbit IgG (H+L)-HRP (111-035-003, Jackson Immune Research, WB: 1:5000); Goat Anti-Mouse IgG (H+L)-HRP (115-035-003, Jackson Immune Research, WB: 1:5000); Streptavidin-HRP (016-030-084, Dianova, WB: 1:5000);

Transfection reagents used in this study were as follows: Lipofectamine LTX & PLUS Reagent (15338100, Thermo Fisher Scientific); Lipofectamine RNAiMAX Reagent (13778150, Thermo Fisher Scientific); PolyFect Transfection Reagent (301105, Qiagen); Nanofectin Kit (Q051-005, PAA Laboratories); and ScreenFect A (299-73203, Wako Laboratory Chemicals).

The following siRNAs were used: siRNA Myo19 (GCAAAUGACUGGAGCCGCA, UAACAACAGCAGUCGCUUU, GGGAGGUCCUGCUGUACAA, CGCCCGAGCUAAUGAGAGA (L-017137-01-0005, Dharmacon, GE Healthcare); siRNA non-targeting (UGGUUUACAUGUCGACUAA, UGGUUUACAUGUUGUGUGA, UGGUUUACAUGUUUUCUGA, UGGUUUACAUGUUUUCCUA (D-001810-10, Dharmacon, GE Healthcare); siRNA Miro-1 (#-09 UGUGGAGUGUUCAGCGAAA, #-10 GCAAUUAGCAGAGGCGUUA, #-11 CCAGAGAGGGAGACACGAA, #-12 GCUUAAUCGUAGCUGCAAA (J-010365-XX, Dharmacon, GE Healthcare); siRNA Miro-2 (#-09 GCGUGGAGUGUUCGGCCAA, #-10 CCUCAAGUUUGGAGCCGUU, #-11 GAGGUUGGGUUCCUGAUUA, #-12 AGGAGAUCCACAAGGCAAA, J-008340-XX, Dharmacon, GE Healthcare); SMARTpool Kif5B siRNA (Dharmacon: ON-TARGETplus KIF5B siRNA L-008867-00-0005).

### Construction of plasmids

Human Myo19 cDNA (AK304073 / FLJ61052 / TRACH 3006685) was obtained from the National Institute of Technology and Evaluation (NITE) Japan. It was subcloned into pEGFP-C1 by PCR amplification of a 5’- and a 3’-fragment, introducing a BglII and an XbaI site, respectively. The plasmid pEGFP-Myo19 tail codes for a fusion protein of EGFP with amino acids 824-970 of Myo19, pEGFP-Myo19 N-tail with amino acid residues 824-897 of Myo19 and pEGFP-Myo19 C-tail with amino acid residues 898-970 of Myo19, respectively. The myc-BirA R118G-MCS (BioID) plasmid was obtained from Addgene (plasmid #35700) (Roux et al., 2012). The Myo19 tail cDNA coding for residues 824-970 was subcloned into the XhoI and HindIII restriction sites. To construct pMyo19-Halo, the Halo-tag flanked by a BshTI restriction site and a stop codon followed by a NotI restriction site was amplified by PCR using plasmid pFN21AA1042 (Kazusa DNA Res. Inst.) as a template. The Halo-tag cDNA was used to replace the cDNA for EGFP in pMyo19-EGFP-N1. pHalo-Myo19 was constructed by replacing EGFP in the plasmid pEGFP-C1-Myo19 with amplified Halo-tag using the restriction sites BshTI and Bgl II. Plasmid pMyo19tail-Halo-N1 was constructed by amplifying Myo19tail from pEGFP-C1-Myo19tail by PCR, introducing BglII and EcoRI restriction sites. This PCR product was then used to replace the Myo19 coding sequence in pMyo19-Halo-N1. Plasmid pHalo-C1-Myo19tail was constructed by amplifying Halo from plasmid pFN21AA1042 by PCR, introducing AgeI and BspEI restriction sites. Using these sites, this PCR product was then used to replace the EGFP coding sequence in pEGFP-C1-Myo19tail-avi-FLAG. To express GST-Myo19 C-tail (aa 898-970)-FLAG, the sequence coding for C-tail-FLAG was amplified by PCR and subcloned into pGEX-4T-1 using EcoRI/SalI restriction sites. pRK5-Myc-Miro-1 and pRK5-myc-Miro-2 plasmids were kind gifts of P. Aspenström, Stockholm, Sweden (Fransson et al., 2003). The plasmid mCherry-Miro-1 was constructed by inserting Miro-1 into the XhoI/SacII sites of pmCherry-C1. The point mutation to create plasmid mCherry-Miro-1 N18 was introduced by QuikChange mutagenesis. The GFP-2xFYVE^Hrs^ plasmid was a kind gift of M. Falasca, London, UK (Maffucci et al., 2003). The plasmid mCherry-Miro-1-ΔTM-2xFYVE has the sequences coding for amino acid residues 592-618 of Miro-1 replaced by the sequences coding for 2xFYVE domains. The plasmid coding for the fusion protein pmCherry-Miro-1-aa1-408-2xFYVE was constructed by standard molecular biology techniques. The pET28a(+) plasmid encoding hMiro1-ABC(aa1-592)-6xHis was a kind gift of S.E. Rice (Chicago, USA) (Klosowiak et al. 2016). The plasmid encoding the mitochondrial targeting sequence from the subunit VIII of the human cytochrome c oxidase fused to RFP was purchased from BD Pharmingen and the sequence coding for RFP was exchanged for EGFP and mCherry, respectively. Plasmid mTagBFP2-TOMM20-N-10 was a gift from Michael Davidson (Addgene plasmid # 55328)(Subach et al., 2011). All fragments amplified by PCR were sequenced.

### Cell culture and transfection of HeLa cells

HeLa cells were cultivated in DMEM supplemented with 10 % (v/v) FCS, 100 U/ml penicillin and 100 µg/ml streptomycin (complete DMEM) at 37 °C, 95 % humidity and 5 % CO_2_.

For transfection of HeLa cells 20,000 cells were seeded onto glass coverslips in 24-well-plates. Cells were transfected one day after seeding as described in the manual of the manufacturer. Briefly, 0.5 µg of the plasmid DNA was mixed with 1 µl Lipofectamine LTX and 0.5 µl PLUS Reagent in 50 µl DMEM, vortexed, incubated at room temperature (RT) for 5 min and added drop wise to the cells. For a cotransfection of two plasmids the amount of plasmid DNA was reduced to 0.25 µg for each plasmid. After 4 h of incubation the DNA-lipid complexes were removed, cells were washed with PBS and fresh complete DMEM was added. Cells were fixed 24 h after transfection.

For live cell imaging, approximately 40,000 HeLa cells were seeded onto a 35 mm µ-dish (ibidi) in 1 ml complete DMEM one day before transfection. For transfection, 2 µg of the plasmid DNA in 50 µl diluent (150 mM NaCl) was mixed with 6.4 µl Nanofectin in 50 µl diluent, and vortexed. The mixture was incubated at RT for 30 min and added to the µ-dish drop wise. The transfection complexes were removed after 4 h of incubation, and fresh complete DMEM was added to the µ-dish. After 24 h of incubation, the expression of recombinant proteins was analyzed.

To generate the stable HeLa myc-BirA* Myo19 tail, Halo-Myo19 and Myo19-Halo cell lines, 100,000 cells per well were seeded in a 6-well-plate. The following day, 2.5 µg of the plasmid DNA with 5 µl Lipofectamine LTX and 2.5 µl PLUS Reagent in 250 µl DMEM was used for transfection. After 48 h complete DMEM was supplemented with 600 µg/ml G418 and 60%, 30% and 10% of cells, respectively, were replated in 150 mm Ø cell culture dishes. Cells still viable after two weeks were considered stably transfected.

Single colonies of cells were isolated with a metal ring, trypsinized and transferred to 6-well-plates. Cells were processed for western blot analysis and further expansion.

For knockdown studies of Miro-1 and Miro-2, 100,000 HeLa cells per well were seeded in 6-well-plates. The following day, cells in one well were transfected with 25 pmol of each siRNA or each pool of 4 siRNAs that was mixed with 7.5 µl Lipofectamine RNAiMAX in 183 µl DMEM. Cells were harvested for western blot analysis 48 h and 72 h after transfection.

### BioID

BioID was carried out essentially as described in Roux et al. (2013) with some modifications. Cells were incubated for 24 h in complete DMEM supplemented with 50 µM biotin before lysis and affinity purification of biotinylated proteins. To enhance expression of the construct, 10 mM Na-butyrate was added to the medium 15 h before harvesting of the cells.

Approx. 2×10^8^ stably transfected myc-BirA* Myo19 tail HeLa cells (10 confluent 150 mm Ø cell culture dishes) were washed three times with cold PBS and collected by scraping. They were resuspended in 20 ml lysis buffer (50 mM Tris, pH 7.4, 500 mM NaCl, 0.2 % (w/v) SDS, 0.1 mg/ml Pefabloc, 0.01 mg/ml Leupetin and 0.02 U/ml Aprotinin) before 2 ml of 20 % (v/v) Triton X-100 were added. The subsequent steps were performed at 4 °C. Lysates were sonicated with two times 30 pulses using an amplitude of 100 % and a proportional sonication period of 0.6 (UP 100H sonicator, Hielscher Ultrasound Technology). Prior to the second sonication cycle 20 ml 50 mM Tris pH 7.4 were added. Lysates were clarified by centrifugation at 16,600 x g for 10 min. Supernatants were incubated with 8 mg of equilibrated streptavidin magnetic beads (Pierce 88817, Thermo Fisher Scientific) overnight. Beads were collected and washed with wash buffer 1 (2% (w/v) SDS in ddH_2_O) for 8 min at RT. This step was repeated with wash buffer 2 (0.1 % (w/v) deoxycholic acid, 1 % (w/v) Triton X-100, 1 mM EDTA, 500 mM NaCl and 50 mM Hepes pH 7.5), wash buffer 3 (0.5 % (w/v) deoxycholic acid, 0.5 % (w/v) NP-40, 1 mM EDTA, 250 mM LiCl and 10 mM Tris pH 7.4) and wash buffer 4 (50 mM Tris pH 7.4), respectively.

Biotinylated proteins were eluted from the beads at 98 °C for 8 min with 200 µl buffer containing 40 mM Tris pH 6.8, 1 mM EDTA, 8 % (w/v) sucrose, 3 % (w/v) SDS, 2 % (w/v) 2-mercaptoethanol, 0.004 % (w/v) bromophenol blue and 820 µM biotin. Aliquots of the eluates were analysed by immunoblotting (20 µl) and by coomassie blue staining following SDS-PAGE (40 µl). Isolated bands were subjected to mass spectrometry (University Hospital Cologne, Centre of Molecular Medicine (ZMM), Central Bioanalytics (ZBA)).

### Expression and purification of recombinant proteins

Miro1 (aa 1-592, C-terminal 6xHis-tag) (Miro1-ABC) was expressed in *E. coli* Rosetta 2 (DE3) cells and purified by metal ion affinity chromatography with TALON beads (Clontech Laboratories, Inc.). Cells were cultured in TPM medium (20 g/L tryptone, 15 g/L yeast extract, 8 g/L NaCl, 2 g/L Na_2_HPO_4_, 1g/L KH_2_PO_4_) containing 50 µg/mL kanamycin and 34 µg/mL chloramphenicol. Protein expression was induced over night at 18 °C with 125 µM IPTG when cell density had reached an OD_600_ of 0.9. Cells were harvested by centrifugation (4,400 x g, 15 min, 4 °C), washed with cold PBS (137 mM NaCl, 2.7 mM KCl, 10.2 mM Na_2_HPO_4_, 1.8 mM KH_2_PO_4_, pH 7.0) and stored at -80 °C. For protein purification, cell pellets were resuspended in lysis buffer (25 mM Hepes, pH 7.4, 300 mM NaCl, 0.5 mM TCEP, 8 mM imidazole, 2 mM MgCl_2_, 5 % sucrose, 0.02 % Tween-20, 0.1 mg/ml Pefabloc®, 0.01 mg/ml Leupetin, 0.02 U/ml Aprotinin, 2 mM ATP) and cells were lysed by sonication for 4 × 2 min with 2 min intervals using an amplitude of 100 % and a proportional sonication period of 0.6 (UP 100H sonicator, Hielscher Ultrasound Technology). The lysate was clarified by ultracentrifugation (139.000 x g, 45 min, 4 °C) and batch adsorbed to preequilibrated TALON beads for 1 h at 4 °C. The beads were collected and washed 3 x with washing buffer (25 mM Hepes, pH 7.4, 300 mM NaCl, 0.5 mM TCEP, 12 mM imidazole, 2 mM MgCl_2_, 5 % sucrose, 0.02 % Tween-20) before they were transferred to a gravity-flow column. Bound protein was eluted with elution buffer (25 mM Hepes, pH 7.4, 300 mM NaCl, 0.5 mM TCEP, 300 mM imidazole, 2 mM MgCl_2_, 5 % sucrose, 0.02 % Tween-20), pooled and stored at -20 °C.

Myo19 C-tail (aa 898-970) fused to a N-terminal GST-tag and a C-terminal FLAG-tag was expressed in *E. coli* Rosetta 2 (DE3) cells and purified by glutathione affinity chromatography followed by FLAG-antibody affinity chromatography. The procedure for cell growth, induction of protein expression and harvest of cells was as described above for Miro1-ABC protein purification. Cell pellets were stored overnight at -80 °C. They were resuspended in lysis buffer (20 mM Hepes, pH 7.4, 100 mM NaCl, 50 mM KCl, 2 mM MgCl_2_, 10 % glycerol, 1 mM 2-mercaptoethanol, 0.1 mg/ml Pefabloc®, 0.01 mg/ml Leupetin, 0.02 U/ml Aprotinin, 2 mM ATP) and lysed by sonication. Following ultracentrifugation (139.000 x g, 45 min, 4 °C), the supernatant was incubated with preequilibrated glutathione sepharose beads. The beads were collected and washed 3 x with washing buffer (20 mM Hepes, pH 7.4, 100 mM NaCl, 50 mM KCl, 2 mM MgCl_2_, 1 mM 2-mercaptoethanol). The protein bound to the resin was eluted by cleavage of the GST-moiety by the addition of 40 U of thrombin overnight in washing buffer. Beads were centrifuged at 700 x g for 5 min at 4°C and washed 2-times. The supernatants were combined and passed several times over FLAG-antibody agarose (ANTI-FLAG® M2 Affinity Gel, Sigma-Aldrich). The column resin was washed with washing buffer, followed by two alternating high salt (20 mM Hepes, pH 7.4, 450 mM NaCl, 50 mM KCl, 2 mM MgCl_2_, 1 mM 2-mercaptoethanol) and low salt (20 mM Hepes, pH 7.4, 50 mM KCl, 2 mM MgCl_2_, 1 mM 2-mercaptoethanol) washes, followed by washing buffer. Finally, Myo19 C-tail was eluted with 0.25mg/mL FLAG®-peptide (Sigma-Aldrich) in washing buffer.

### Pull-down assay

Purified hMiro1-ABC was loaded with GTP by adding 2 mM GTP and 6 mM EDTA for 10 min at RT followed by the addition of 15 mM MgCl_2_. Next, the protein was diluted with assay buffer (25 mM Hepes, pH 7.4, 300 mM NaCl, 0.5 mM TCEP, 2 mM MgCl_2_, 5 % sucrose, 0.02 % Tween-20) to a final concentration of 10 mM imidazole and adsorbed to preequilibrated TALON beads at 4 °C. Beads were washed three times with washing buffer (25 mM Hepes, pH 7.4, 300 mM NaCl, 0.5 mM TCEP, 12 mM imidazole, 2 mM MgCl_2_, 5 % sucrose, 0.02 % Tween-20) and transferred to a gravity-flow column. Myo 19 C-tail was passed repeatedly over the column before it was washed three times with washing buffer. Adsorbed Miro1-ABC was eluted with an excess of imidazole in the buffer (25 mM Hepes, pH 7.4, 300 mM NaCl, 0.5 mM TCEP, 400 mM imidazole, 2 mM MgCl_2_, 5 % sucrose, 0.02 % Tween-20). Eluted proteins were analyzed by SDS-PAGE and immunoblotting.

### Preparation of cell homogenates

Cells grown in 6-well-plates were washed three times with cold PBS and gently scraped off. They were centrifuged at 600 x g at 4 °C for 5 min and resuspended in 100 µl lysis buffer (50 mM Tris pH 7.4, 2 mM MgCl_2_, 100 mM NaCl, 1 % (v/v) NP-40, 10 % (v/v) glycerol, 1 mM DTT, 0.1 mg/ml Pefabloc, 0.01 mg/ml Leupeptin and 0.02 U/ml Aprotinin). Protein concentrations were determined by Bradford assay using BSA as a standard. Equal amounts of protein were separated by SDS-PAGE and further processed for immunoblotting.

### Pulse-chase experiments

For pulse-chase experiments stable Halo-Myo19 or Myo19-Halo HeLa cells were transfected with siRNA for Miro1+2. 24h later, the cells were pulse labeled with 2 µM HaloTag® TMR ligand (Promega) for 30 min at 37°C, 97 % humidity and 5 % CO_2_. Free ligand was removed by several washing steps and a 30 min incubation with complete DMEM supplemented with 600 µg/ml G418. Next, the cells were incubated with 5 µM HaloTag® Biotin ligand, to prevent further labeling of the Halo-tag with any remaining fluorescent ligand. Cells were harvested at 0, 4, 8 and 12 h time points. Proteins were separated by SDS-PAGE, electrophoretically transferred to a PVDF membrane and analyzed for TMR fluorescence followed by immunoblotting. Fluorescent bands were quantified using Image J.

### Mitochondria purification

Prior to mitochondria purification, HeLa wt and HeLa myc-BirA* Myo19 tail cells were treated overnight with 10 mM Na-butyrate. Mitochondria were purified with Qproteome Mitochondria Isolation Kit (QIAGEN) according to the manufacturers protocol for high-purity mitochondria.

### Fluorescence stainings

To label mitochondria, cells were incubated with 75 nM MitoTracker Red CMXRos (M7512, Thermo Fisher Scientific) in complete DMEM at 37 °C, 95 % humidity and 5 % CO_2_ for 10 min. Immediately afterwards, cells were washed three times with warm PBS and fixed with 4 % (w/v) PFA in PBS at 37 °C for 25 min. Free aldehydes were quenched with 0.1 M glycine in PBS at RT for 10 min. Subsequently, cells were washed three times with PBS and mounted with Mowiol (3.4 mM Mowiol 4-88, 105 mM Tris pH 8.5, 18.4 % (v/v) Glycerin and 223 mM DABCO). To stain cells for endogenous Myo19, they were permeabilized at RT for 15 min with 0.1 % (v/v) Triton X-100 in PBS, followed by washing with PBS for 2x 5 min. Unspecific binding sites were blocked by incubation with blocking buffer (5 % (v/v) normal goat serum in PBS) at RT for 10 min. The primary antibody (HPA021415, Sigma-Aldrich) was diluted 1:100 in blocking buffer. Coverslips were placed on a drop of antibody solution in a humid chamber and incubated overnight at 4 °C. Thereafter, cells were washed 3x 5 min with PBS at RT and incubated in the dark with secondary antibody (111-545-003, Jackson ImmunoResearch) diluted 1:500 in blocking solution for no longer than 1 h at RT. Subsequently, cells were washed three times with PBS and mounted with Mowiol (3.4 mM Mowiol 4-88, 105 mM Tris pH 8.5, 18.4 % (v/v) Glycerin, 223 mM DABCO).

### Transferrin endocytosis assay

HeLa cells grown on coverslips were serum-starved for 30 min in MEM medium at 37 °C, 5 % CO_2_ and 95 % humidity. AF 488-conjugated human transferrin (30 µg/ml) (ThermoFisher) in MEM medium was added for 1 h at 37 °C. Cells were washed with ice cold 1x PBS and subsequently fixed with 4 % PFA for 20 min at RT.

### Retargeting and quantification

HeLa cells were fixed 24 h after transfection and analysed by spinning disk microscopy (UltraVIEW VoX, PerkinElmer; Nikon Eclipse Ti). Images were acquired with a 60x/1.49 NA oil objective and an EMCCD camera using Volocity software. Z-stacks covering the entire depth of cells with intervals of 0,5 µm were acquired. To quantify retargeting of tail constructs to early endosomes maximum intensity projections were obtained using ImageJ. To separate mitochondria from background a “top hat” filter was applied.

Pearson’s correlation coefficient was used to express the intensity correlation of co-localizing objects. Analysis was restricted to custom ROI of each cell, with a thresholding factor of 2. The result is +1 for perfect correlation, 0 for no correlation, and -1 for perfect anti-correlation.

### Mitochondria distribution

#### Imaging

To assess mitochondria distribution, 20’000 HeLa cells per well were plated on glass coverslips in 24-well plates. The next day, cells were transfected with appropriate plasmids using the transfection reagent Lipofectamine^TM^ LTX and PLUS^TM^ reagent (Invitrogen) according to the manufacturer’s instructions. 24 h after transfection, the cells were processed further: Cells that had been transfected with Halo constructs were labelled with 5 µM HaloTag® TMR ligand (Promega) in full medium for 30 min at 37 °C, followed by several washing steps with PBS and full medium before fixation. Cells were washed with PBS and fixed with 4% PFA, stained for DNA with DAPI and, if appropriate, for F-actin with fluorophore-labelled phalloidin, mounted on microscope slides and left to harden at 4 °C overnight. For experiments involving protein depletion, 100’000 HeLa cells per well were seeded in 6-well plates. 24 h after seeding, the cells were transfected with appropriate siRNAs using the transfection reagent Lipofectamine^TM^ RNAiMAX according to the manufacturer’s instructions. The next day, the cells were replated into 24 well plates and transfected, fixed and stained as described above. Imaging was performed with a 63x N.A. 1.4 Plan-Apochromat oil immersion objective on a Zeiss LSM 510 confocal laser scanning microscope equipped with Argon and Helium-Neon lasers and three 12-bit R/FI detectors. Z-stacks of mitochondria and, if appropriate, other labeled structures were recorded at a plane interval of 0.5 µm; upper and lower boundaries of the stacks were set manually for each field of view. Bright-field images were recorded with 12-bit R/FI detectors along with the z-stacks. DAPI fluorescence was excited with a mercury vapor short-arc HBO 50 lamp and detected with an 8-bit AxioCam MRm camera.

### Analysis

Image processing and analysis were performed in ImageJ (NIH, USA) with customized macros. Briefly, z-stacks of fluorescence channels were converted to sum projections, and sum projections of the mitochondria were top-hat filtered and background corrected. Size and orientation of the DAPI images were adjusted to the confocal images. For each cell, the outline was determined manually based on the bright-field image. Shape descriptors routinely available in ImageJ were measured for the cell outline. The nucleus outline was determined via automated thresholding of the DAPI image and was dilated to determine the boundary of the perinuclear region, which was defined as a 4 µm wide ring around the nucleus. When the dilated nucleus outline exceeded the cell outline, it was cut accordingly. For each of the cell’s regions (in/above/below nucleus, perinuclear, peripheric), the proportion of mitochondria signal was measured as well as the fraction of the total cell area occupied by this region. The perinuclear mitochondria accumulation factor was defined as the proportion of mitochondrial signal within the perinuclear region, divided by the fraction of the total cell area occupied by the perinuclear region.

### Statistical evaluation

Data are displayed as box plots with the box ranging from 25^th^ to 75^th^ percentiles, whiskers ranging from 10^th^ to 90^th^ percentiles, the median marked as a line, and the mean marked as a square. Data were analyzed by non-parametric statistical tests; the significance levels were set to p ≤ 0.05 (*), p ≤ 0.01 (**), and p ≤ 0.001 (***).

## ACKNOWLEDGMENTS

This work was supported by the DFG (BA 1354/10-1). K.M was supported by the Cells in Motion Excellence Cluster (CiM) / International Max-Planck Research School (IMPRS) graduate school and X-P.H. by the NRW Research School "Cell Dynamics and Disease, CEDAD". S.J.O. acknowledges the receipt of a CiM bridging position. We are grateful to P. Aspenström, M. Falasca and S.E. Rice for the gift of plasmids, M. Horsthemke for help with artwork and to P. Hanley for valuable comments on the manuscript.

---

**Supplementary Figure S1: Schematic representations of Myo19 and Miro-1 constructs.** A) Schematic representation of human Myo19 and of the tail constructs that were used in the study. The three bars colored brown indicate the three IQ-motifs that form the light chain binding domain which separates head and tail regions of Myo19. B) Schematic representation of human Miro-1 constructs. TM, transmembrane domain; ELM, EF-hand pair with ligand mimic. Fluorescent proteins (not indicated) were fused to the N-terminus of all constructs in A) and B). Numbers indicate amino acid residues.

**Supplementary Figure S2: Overexpression of Myo19 tail induces the loss of endogenous Myo19.** Wild-type (wt) and stably with myc-BirA*-Myo19 tail (tail) transfected HeLa cells were incubated overnight without (-) or with (+) Na-butyrate to enhance expression of the recombinant protein. Proteins of cell homogenates were separated by SDS-PAGE and detected by immunoblotting. Na-butyrate enhanced the expression of myc-BirA*-Myo19 tail which was accompanied by reduced levels of endogenous Myo19.

**Supplementary Figure S3: Miro is retargeted to early endosomes and this retargeting is independent of Myo19 tail constructs.** A) HeLa cells were transfected to express mCherry-Miro1 that had its C-terminal transmembrane domain replaced by 2xFYVE domains (EE) which bind to early endosomes (Miro1 ΔTM-EE). These cells were incubated with fluorescent transferrin (Transferrin 488) to label the endocytic pathway and imaged. Miro1 ΔTM-EE was co-localized with some of the endosomes labeled by transferrin. B) Different mCherry-Miro1 constructs (Miro1 ΔTM-EE, Miro1 aa1-408-EE and Miro1 T18N ΔTM-EE) targeted to early endosomes demonstrate only a marginal colocalization with mitochondria (TOMM20) independent of the co-expressed Myo19 tail constructs (Tail, N-tail, C-tail). Analysis was performed with triple transfected cells; n≥30 from three independent experiments.

**Supplementary Figure S4: Overexpression of full length EGFP-Myo19, but not of Myo19 fragments lacking the motor domain, impacts the size and shape of cells.** HeLa cells overexpressing GFP-Myo19 covered smaller areas on the substratum, had shorter perimeters and were rounder than GFP-overexpressing control cells. Myo19 fragments lacking the motor domain did not cause these changes. n = 153-159 cells per construct from 3 independent experiments.

